# Fibronectin: a natural barrier against prion infection

**DOI:** 10.1101/2023.08.31.555658

**Authors:** M Carmen Garza, Sang Gyun Kang, Chiye Kim, Eva Monleón, Jacques van der Merwe, David A. Kramer, Richard Fahlman, Valerie Sim, Judd Aiken, Debbie McKenzie, Leonardo M. Cortez, Holger Wille

**Affiliations:** Centre for Prions and Protein Folding Diseases, Neuroscience and Mental Health Institute; Departments of 2Biochemistry; Biological Sciences; Laboratory Medicine and Pathology; Medicine; Neuroscience and Mental Health Institute; Agriculture, Food and Nutritional Science, University of Alberta, Edmonton, Alberta; Centro de Encefalopatías y Enfermedades Transmisibles Emergentes, Departamento de Anatomía e Histología Humana, Universidad de Zaragoza, IA2, IIS Aragón, Zaragoza, Spain

**Keywords:** Fibronectin, extracellular matrix (ECM), reserve cells, myotube, muscle, prion, scrapie, tropism

## Abstract

A distinctive signature of the prion diseases is the accumulation of the pathogenic isoform of the prion protein, PrP^Sc^, in the central nervous system of prion-affected humans and animals. PrP^Sc^ is also found in peripheral tissues, raising concerns about the potential transmission of pathogenic prions through human food supplies and posing a significant risk to public health. Although muscle tissues are considered to contain levels of low prion infectivity, it has been shown that myotubes in culture efficiently propagate PrP^Sc^. Given the high consumption of muscle tissue, it is important to understand what factors could influence the establishment of a prion infection in muscle tissue. Here we used *in vitro* myotube cultures, differentiated from the C2C12 myoblast cell line (dC2C12), to identify factors affecting prion replication. A range of experimental conditions revealed that PrP^Sc^ is tightly associated with proteins found in the systemic extracellular matrix (ECM), mostly fibronectin (FN). The interaction of PrP^Sc^ with FN decreased prion infectivity, as determined by standard scrapie cell assay. Interestingly, the prion-resistant reserve cells in dC2C12 cultures displayed a FN-rich ECM while the prion-susceptible myotubes expressed FN at a low level. In agreement with the *in vitro* results, immunohistopathological analyses of tissues from sheep infected with natural scrapie demonstrated a prion susceptibility phenotype linked to an extracellular matrix with undetectable levels of FN. Conversely, PrP^Sc^ deposits were not observed in tissues expressing FN. These data indicate that extracellular FN may act as a natural barrier against prion replication and that the extracellular matrix composition may be a crucial feature determining prion tropism in different tissues.

**Author summary:** Prion diseases are complex fatal neurodegenerative disorders caused by a misfolded form of the cellular protein PrP^C^ (PrP^Sc^). Due to the potential zoonotic transmission of these disorders through animal-based food intake, it is crucial to identify the tissues in which PrP^Sc^ can accumulate and what might influence its tropism. Animal muscle (or meat) and related food products are highly consumed, raising concern of the involvement of muscle cells in prion replication. Muscle tissue from prion-affected animals contains low levels of infectivity and PrP^Sc^ is mostly associated with nerve structures rather than myofibers, whereas C2C12 myotubes, a muscle-derived cell type, efficiently replicate prions *in vitro* and generate high levels of infectivity compared with other cell cultures. We demonstrate a fibronectin-mediated interference with prion infection in differentiated C2C12 cultures that correlates with the findings in tissues from naturally scrapie-infected animals. Our results suggest that extracellular matrix composition, specifically regarding the presence of fibronectin, might determine prion tropism and dissemination.

## Introduction

Prion diseases are fatal neurodegenerative disorders that include Creutzfeldt-Jakob disease (CJD) in humans, bovine spongiform encephalopathy (BSE) in cattle, scrapie in sheep, and chronic wasting disease (CWD) in cervids. These diseases are associated with the conversion of the host-encoded prion protein, PrP^C^, into a misfolded aggregated form, PrP^Sc^ [1]. PrP^Sc^ accumulates preferentially in the central nervous system (CNS), but detectable amounts of PrP^Sc^ can also be found in peripheral tissues of prion-infected animals [2,3]. Lymphoid tissue and the peripheral nervous system are the major sites involved [4–7], but PrP^Sc^ is also present in muscle from sheep with scrapie and cervids with CWD [8–10]. Muscle is of particular concern, given the high rate of human consumption for this type of tissue. Several studies of experimental and natural prion infection have concluded that muscle can be classified as a low infectivity tissue [11], with 5,000-fold less infectivity than brain in case of scrapie-infected sheep [8,9]. The lower levels of infectivity in muscle and the fact that scrapie and CWD are considered non-zoonotic prion diseases resulted in the exclusion of muscle from specific risk materials seized at the slaughterhouse.

The accumulation of PrP^Sc^ in myocytes of both terminal and preclinical stages of experimentally infected animals has been reported, but detailed histopathological studies demonstrated that PrP^Sc^ was mostly associated with peripheral nerves or neuromuscular junctions rather than with myofibers [12,13]. In natural scrapie, we reported that PrP^Sc^ associated with intramuscular nerve fibers and neuromuscular spindles [10]. The same affected nerve structures have been described in experimental scrapie [9], human prion diseases [14,15], experimental infections with BSE [16,17] and hamsters inoculated with transmissible mink encephalopathy [18]. Thus, *in vivo* experiments have not shown clear prion deposition in myofibers of striated muscle. The factors that contribute to the inefficiency of muscle cells as a replication and accumulation site for prions remain unknown.

Interestingly, C2C12 myotubes, a muscle-derived cell type that retains most of the features of myofibers, support prion propagation with ten times higher yield than other mammalian cell lines, including the neuroblastoma N2a cell line [19]. Although the lack of cell division of myotubes is the most logical explanation for the elevated levels of prion accumulation compared to N2a cells, the highly efficient prion replication in myotubes is surprising considering the low susceptibility of muscle cells to prion infection *in vivo*. C2C12 is a murine myoblast cell line that undergoes terminal differentiation to myotubes upon serum removal. In addition to myotubes, a subset of cells, called reserve cells, remains undifferentiated [20]. Contrary to the high susceptibility of myotubes, reserve cells and myoblasts are resistant to prion infection [21,22].

Here, we used a differentiated C2C12 cell line (dC2C12) to explore the factors that influence susceptibility/resistance in these two muscle cell types. We then evaluated if the same factors are involved in the low susceptibility of muscle and other peripheral tissues to prion infection *in vivo*. Our studies indicate that proteins from the extracellular matrix (ECM) play a central role in prion distribution. The negative correlation between PrP^Sc^ deposition and fibronectin (FN) expression observed in dC2C12 reserve cells was also confirmed in different peripheral tissues of naturally infected animals.

## Results

### 1. PrP^Sc^ replicates and accumulates efficiently in myotubes but not in reserve cells

Following serum deprivation, murine-derived C2C12 myoblast cells are terminally differentiated into mono-nucleated reserve cells and multinucleated myotubes [23]. After four days of differentiation, when at least one complete layer of myotubes has formed with reserve cells intermingled (S1 Fig), the cultures were exposed to the murine prion strain, RML, or to normal brain homogenate (NBH). In previous studies, *de novo* generated PrP^Sc^ was detected from day 15 post-infection by immunoblotting [19]; PrP^Sc^ were, by immunocytochemistry (ICC), associated with multi-nucleated cells but not with mononucleated cells [21]. Here we examined the entire dC2C12 culture, at 20 days post-infection, through serial scans along the z-axis with double staining ICC for the prion protein and desmin, a major intermediate filament protein present in muscle-derived cells (Fig 1). PrP^Sc^ was distributed throughout the multilayered culture and appears as a cytoplasmic-fine or coarse-granular staining pattern exclusively associated with myotubes, that range in width from 5 μm to > 56 μm. Prion infection of dC2C12 influenced cell morphology, with a higher proportion of thin myotubes in RML-infected than in NBH-treated cultures (S2 Fig). In contrast to myotubes, all reserve cells were negative for PrP^Sc^, not only those located at the bottom layers of the culture but also those in close contact with PrP^Sc^ positive myotubes, confirming the different susceptibility of these two types of cells to prion infection.

**Fig 1.**
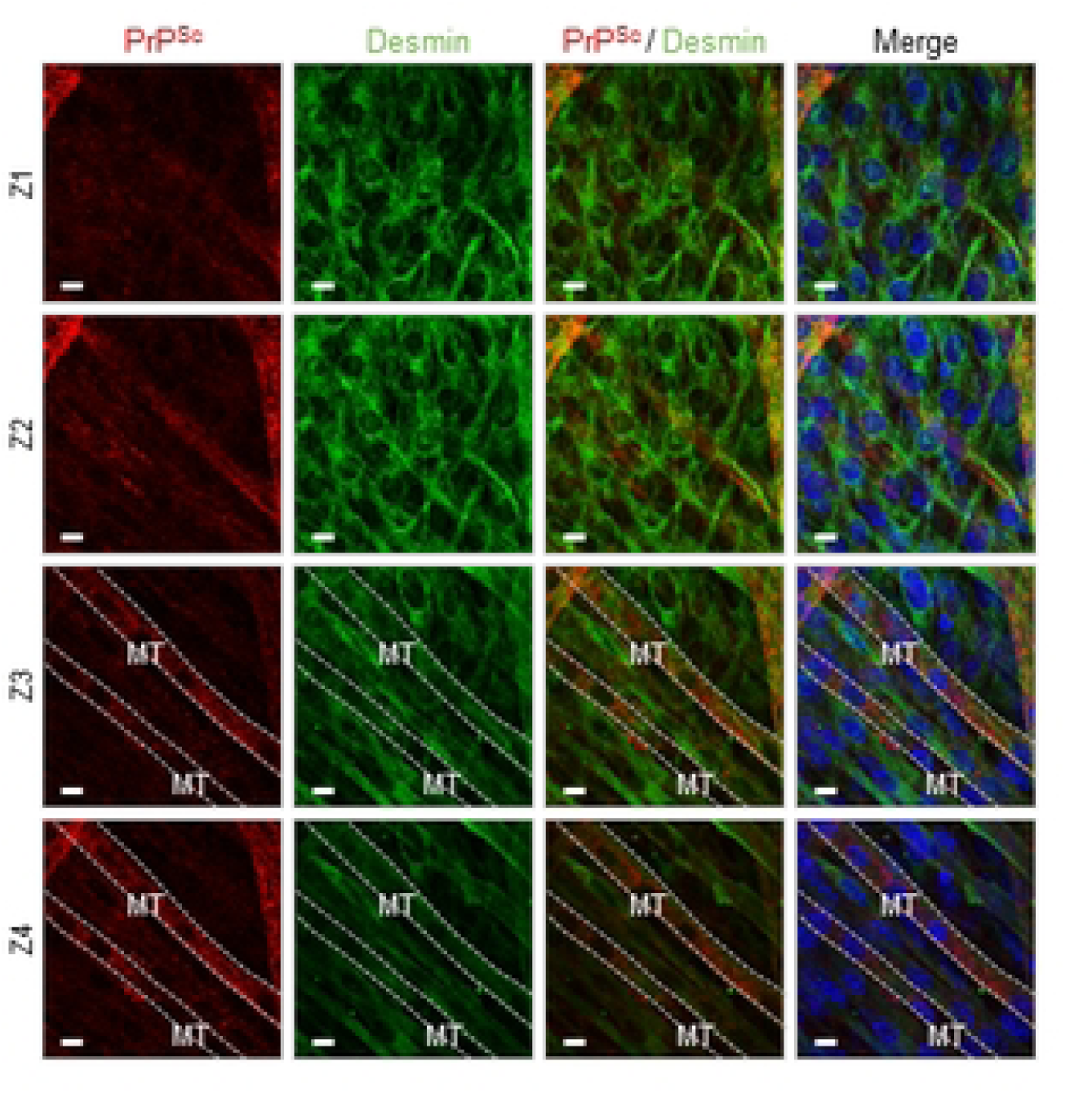
Prion deposition is associated with myotubes in RML-infected dC2C12 cultures at 20 days p.i. Cultures were immunohistochemically stained with SAF83 anti-PrP Ab (red), anti-desmin Ab (green) and DAPI (blue). Four serial micrographs taken along the Z-axis show the multilayer culture composed of 2 morphologically different cell types, the long multinucleated myotubes (MT) abundant in the upper layers (Z3, Z4) and the mononucleated spindle-shape reserve cells (RC), mostly in the bottom layers (Z1, Z2). PrP^Sc^ immunostaining appears as fine intracytoplasmic punctae solely in myotubes with no PrP^Sc^ deposits found in reserve cells. Interval of the confocal Z-stack images is 0.5 µm. Scale bar, 20 µm.

### 2. High molecular weight proteins co-purify with dC2C12 PrP^Sc^ and reduce its infectivity

Uninfected myotubes and reserve cells showed similar levels of PrP^C^ by immunoblotting [19], excluding the possibility that PrP^C^ abundance is the primary factor influencing PrP^Sc^ accumulation in dC2C12.

To investigate whether a cellular component of reserve cells could be interfering/inhibiting PrP^Sc^ replication, we compared infectivity of whole dC2C12 lysate (consisting of PrP^Sc^ from myotubes and any potential inhibitory factors from reserve cells) to PrP^Sc^ purified from the same lysate.

To obtain highly enriched PrP^Sc^ from dC2C12 cultures, we applied a protocol based on protease digestion of the cell lysate followed by selective precipitation of PrP^Sc^ with phosphotungstic acid (PTA) in the presence of sarkosyl [24]. This PTA protocol yielded highly enriched PrP^Sc^ when applied to RML brain homogenate (Fig 2, lane#1). Western blot analysis of the purified fraction showed the three characteristic PrP^Sc^ bands corresponding to di-, mono-, and un-glycosylated PrP isoforms. Silver staining indicated a purity > 95% for PrP^Sc^ in this preparation. Interestingly, much lower purity was obtained when the protocol was applied to dC2C12 lysate (Fig 2, lanes#3-16). Although PrP^Sc^ was detected by immunoblotting, high levels of high molecular weight (HMW) proteins (>100 kDa) co-purified with PrP^Sc^ (Fig 2, lanes #3-16, silver staining). We tested the effect of different detergents in the lysis buffer on increasing PrP^Sc^ yield and decreasing contaminating HMW proteins. Inclusion of sarkosyl yielded the highest amount of PrP^Sc^ although the amount of contaminant protein was also the highest for this condition (Fig 2, lanes#8 and 14). The use of triton X100 - deoxycholate (TD) buffer resulted in a partial reduction of the contaminating proteins but to the detriment of the PrP^Sc^ yield (Fig 2, lanes#9 and 15). RIPA buffer yielded similar levels of PrP^Sc^ as TD but with higher levels of contaminant (Fig 2, lanes#7 and 13). These co-precipitated HMW proteins were also highly resistant to protease digestion (100 - 1000µg/ml of proteinase K, 100µg/ml of pronase E and collagenase). Other conditions such as mild to hard clarification spins before or after benzonase treatments, different centrifugation speeds, stringent sonication), precipitation with different concentrations of PTA and high salt concentration extraction, were tested without a significant increase in purity or yield. Aliquots of the RML brain homogenate used to infect dC2C12 culture were processed in parallel in all conditions showing no HMW proteins contamination for any of them (treatment with sarkosyl and 100µg/ml of PK is shown in Fig 2, lane#1).

**Fig 2.**
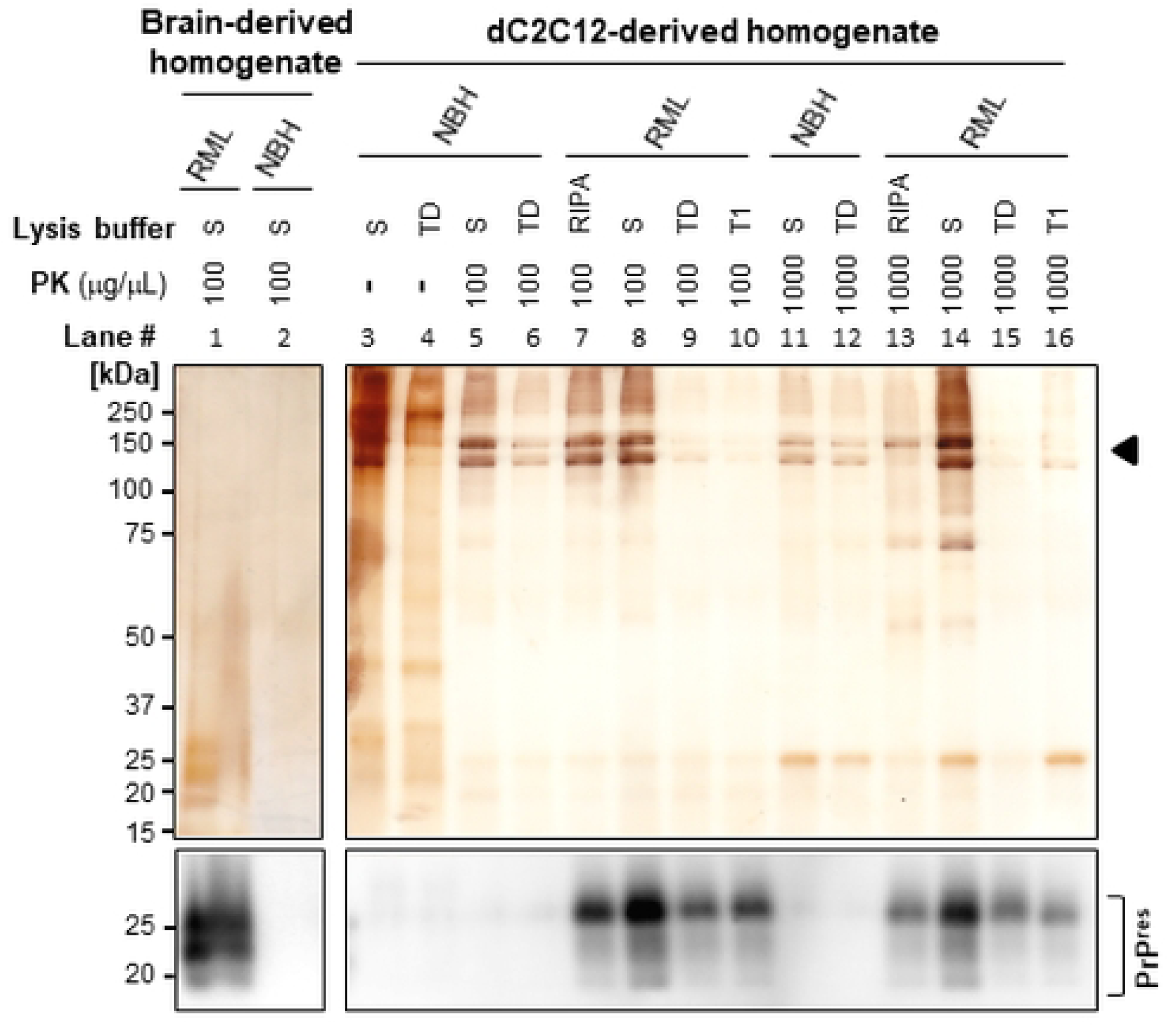
Silver stain and immunoblot analysis of PTA-purified PrP^Sc^ from RML-infected mouse brain and dC2C12 cultures. NBH and RML-infected PTA precipitated PrP^Sc^ samples were lysed with sarkosyl (S), triton deoxycholate (TD) or RIPA buffer, digested with different concentrations of PK and analyzed by SDS-PAGE followed by silver staining (top) or immunoblotting (bottom). High molecular weight (HMW) contaminating proteins >100 kDa (arrow head) were found under all purification conditions for dC2C12 samples. SAF83 dil. 1:10,000 was used to detect PrP.

We then evaluated the infectivity of the whole dC2C12 lysate (in TD lysis buffer) and its PTA-purified PrP^Sc^ (Fig 2, lane#9) by standard scrapie cell assay (SSCA) [25] (Fig 4A). We used PrP^Sc^ purified from the identical volume of dC2C12 lysate that was used in the whole lysate SSCA experiment. The infectivity of PTA-purified PrP^Sc^ was significantly higher (439.3 +/- 125.3 spots/20,000 cells; ave +/- SD) than the infectivity of the whole dC2C12 lysate (283.6 +/- 62.5 spots/20,000 cells; ave +/- SD) suggesting the presence of inhibitory factor/s affecting prion replication in dC2C12 cell cultures.

To evaluate the influence of the HMW contaminating proteins on PrP^Sc^ infectivity, we applied velocity gradient centrifugation to separate these particles based on their size. Given that detergents influence the aggregation state of the PrP^Sc^, and thus their mobility in the gradient [26], we treated infected dC2C12 cells and brain homogenate (BH) with sarkosyl or TD lysis buffers. The solubilized samples were PK digested, loaded on top of an iodixanol gradient and, after centrifugation, fourteen fractions were collected and analyzed by SDS-PAGE, silver staining and immunoblotting (Fig 3). In agreement with Tixador *et al*. [26], the detergent used for solubilization of RML-infected BH influenced migration of PK resistant PrP (PrP^res^), with the PrP^res^ signal in the top half of the gradient when sarkosyl was used (Fig 3A, left) and in the bottom half when the sample was solubilized with TD (Fig 3A, right). Unlike PrP^res^, the HMW proteins were present in the top fractions regardless of the detergent used, but very little was found in BH compared to the dC2C12 samples. Thus, TD solubilized samples yielded highly pure PrP^res^ in fractions #9-13 (Fig 3A right).

**Fig 3.**
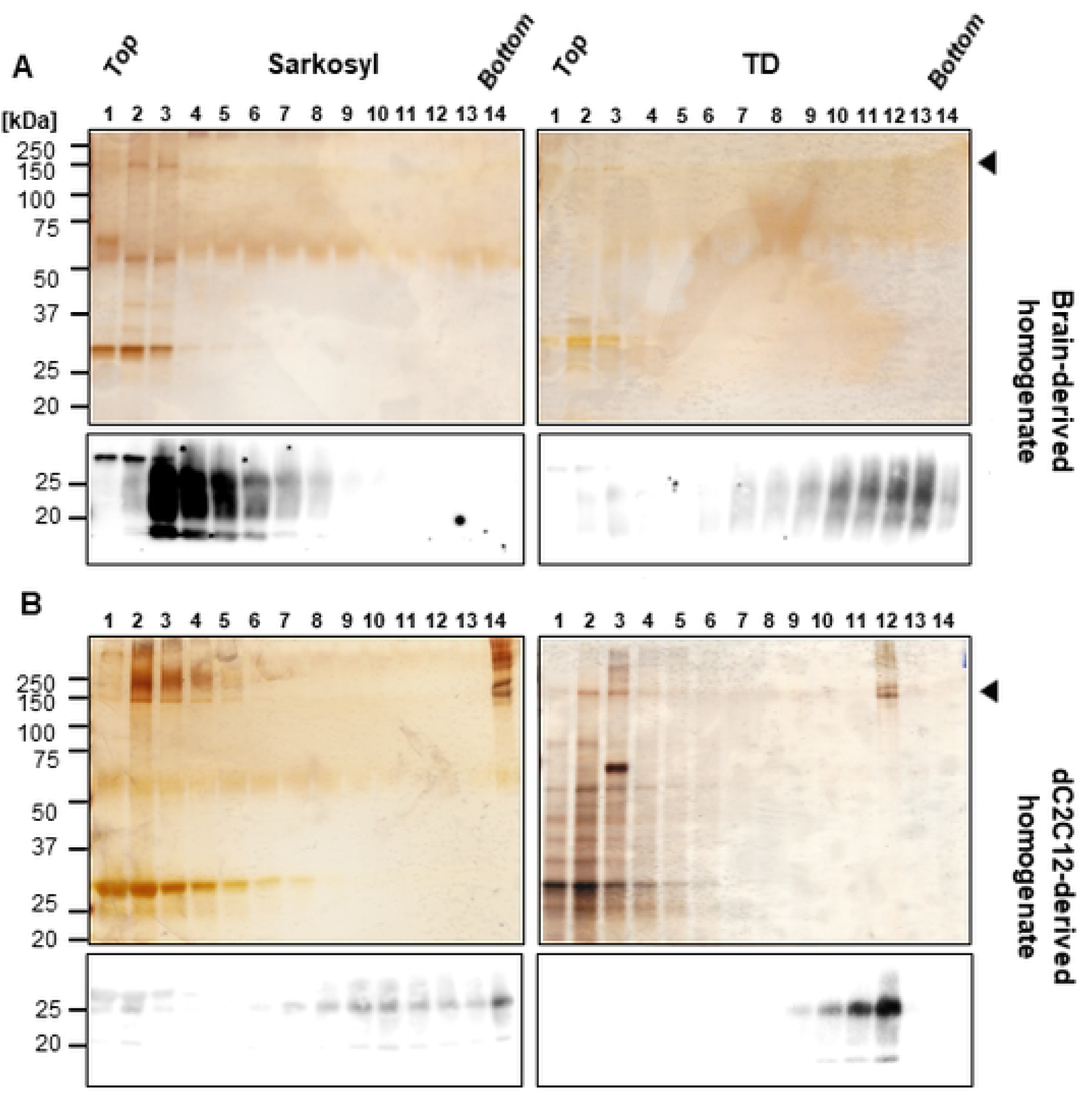
Silver staining and immunoblotting of fractions from velocity gradients. RML-infected mouse brain homogenate (A) and dC2C12 culture (B) were lysed in sarkosyl (left) or TD (right) buffer, PK digested, and the proteins were fractionated by velocity gradient centrifugation in an iodixanol gradient. The presence of HMW proteins (arrow head) was evaluated by silver staining and PrP^Sc^ by immunoblotting (SAF83, dil. 1:10,000) in the 14 fractions collected from top to bottom.

Surprisingly, PrP^res^ from dC2C12 cultures migrated in the bottom half of the gradient with both detergents (Fig 3B). In sarkosyl-solubilized dC2C12 samples, PrP^res^ was distributed more broadly, from fractions #7 to #14, along the gradient, whereas, in the TD solubilized samples, PrP^res^ was detected in fractions #9 to #12. Interestingly, in both conditions, the fraction with higher PrP^res^ signal (fraction #14 for sarkosyl and fraction #12 for TD) also had the highest amount of HMW contaminant proteins, suggesting a physical interaction between these two proteins. In addition, fractions #9 to #11 for sarkosyl and fractions #10 to #11 for TD, contained highly pure PrP^res^. The evaluation of the fractions by SSCA revealed an inhibitory effect of the co-migrating HMW proteins on PrP^Sc^ infectivity (Fig 4B). The TD-treated samples showed that fraction #11, which contained pure PrP^res^, infected the L929 cells with 320 spots/20,000 cells. However, the subsequent fraction, #12, with a higher concentration of PrP^res^ and most of the HMW proteins, was very inefficient at infecting the L929 cells, with fewer spots than the threshold of 150 spots/20,000 cells, indicating an inhibitory effect of the HMW proteins in prion replication. The experiment was performed in triplicate confirming these results. None of the fractions from the sarkosyl-solubilized sample showed infectivity in SSCA.

**Fig 4.**
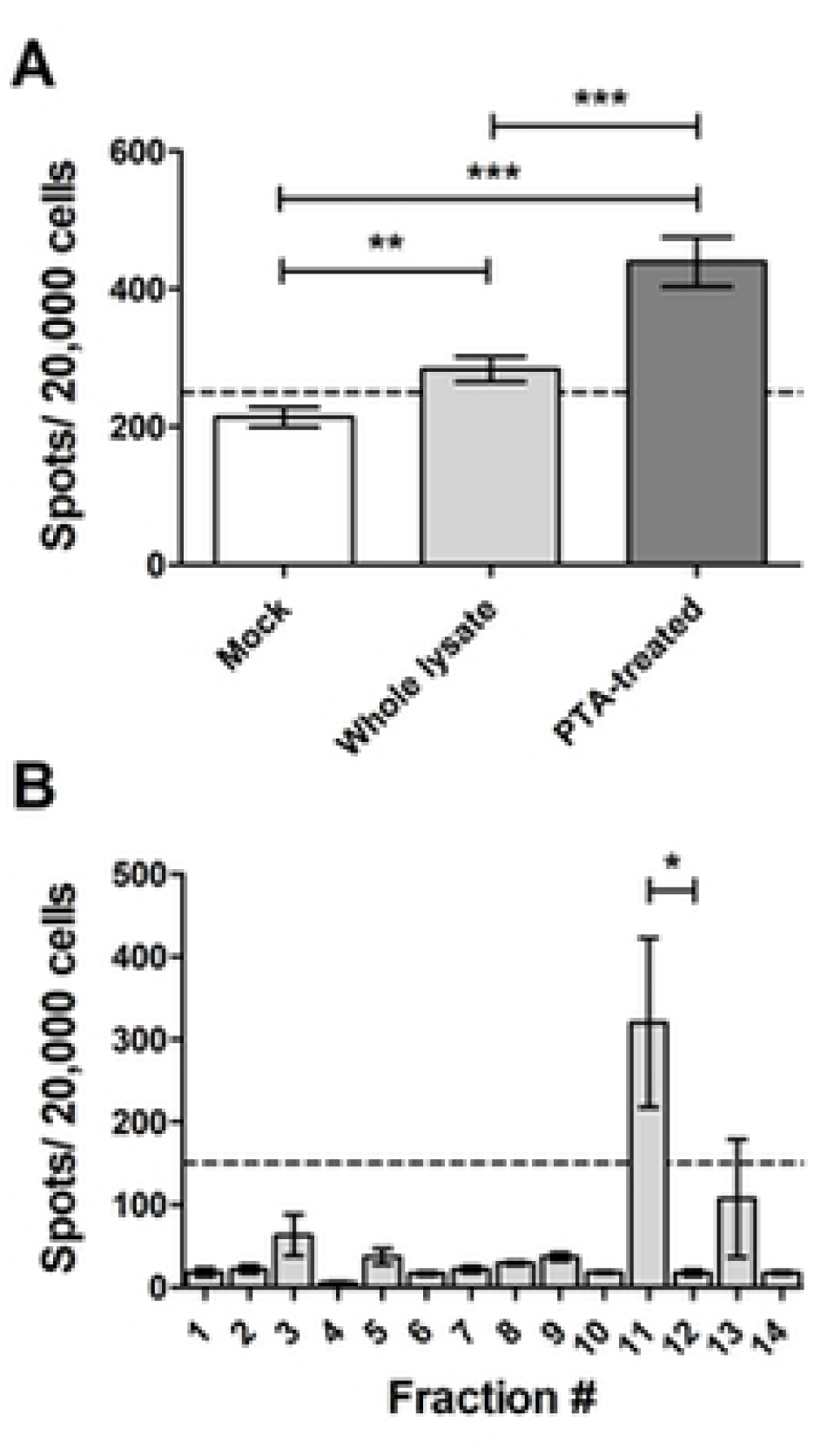
Infectivity of PrP^Sc^ from dC2C12 before and after purification evaluated by SSCA. (A) SSCA of whole lysate (light grey bar) and PTA-purified PrP^Sc^ (dark grey bar) from the same RML infected dC2C12 culture. An uninfected dC2C12 lysate (NBH, white bar) was also analyzed. (B) SSCA of the RML-infected dC2C12 fractions collected after velocity gradient centrifugation (compare Fig 3B). In all cases, dC2C12 cells were lysed in TD buffer. Dotted line represents the cut off threshold for prion infection. Average ± SE was plotted.

### 3. PrP^Sc^ from dC2C12 interacts with extracellular matrix proteins

We next identified the HMW proteins that co-precipitated and co-migrated with PrP^res^ and inhibited its infectivity. The bands larger than 100 kDa detected by silver staining after PTA precipitation or gradient centrifugation were analyzed by mass spectrometry (MS). Fibronectin (FN), collagen and fibrillin, three proteins typically present in the extracellular compartment, were the most prevalent proteins. Table 1 shows the HMW proteins identified by MS in two fractions shown in figure 3. The MS results were validated by immunoblotting (Fig 5A).

**Fig 5.**
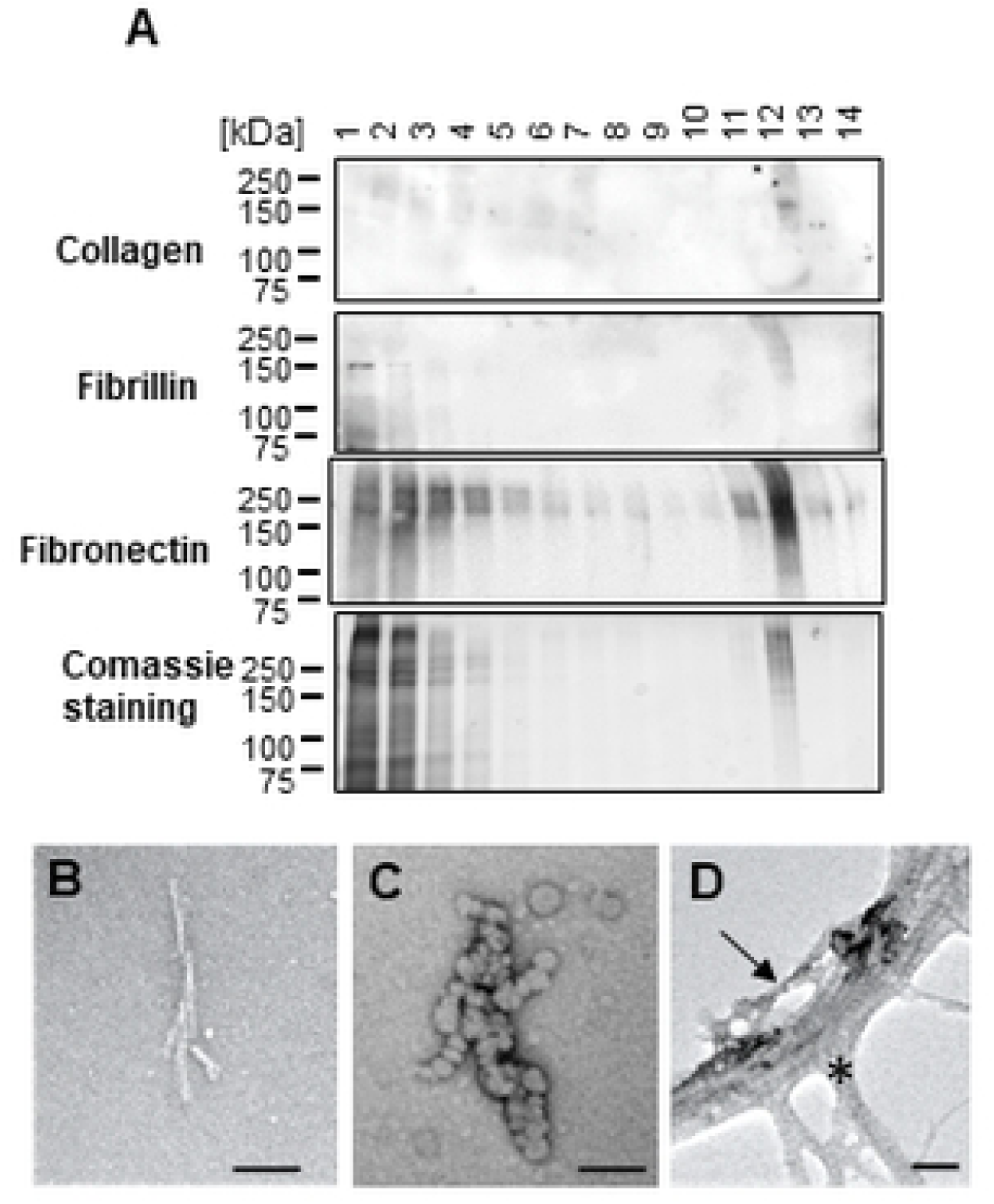
Confirmation of ECM proteins by immunoblotting and presence of prion-like rods confirmed by EM. (A) Immunobloting of collagen, fibrillin and fibronectin and Coomassie blue staining for the fractions of RM-infected dC2C12 cells after velocity gradient centrifugation. (B, C). Fractions from velocity gradients with strong PrP^res^ signal but no HMW bands (following Coomassie staining), were examined by EM. Prion rods were detected in cells lysed with Sarkosyl (C) whereas, following triton X-100 extraction, amorphous aggregates like blobs and oligomers were observed (B). EM examination of a PTA-precipitated sample from RML-infected dC2C12 cultures contains a mesh of fibrillar structures, such as collagen (asterisk), in which a prion rod-like structure (arrow) can be distinguished. Scale bar 50nm.

**Table 1.** HMW proteins detected by MS.

**Fraction 14 of SV gradient from C2C12 in Sarkosyl (Figure 3)**
| Accession | Description | Percentage of Contamination |
| --- | --- | --- |
| P11087 | Collagen alpha-1(I) chain (GN=COL1A1) | 75.6 |
| Q01149 | Collagen alpha-2(I) chain (GN=COL1A2) | 11.9 |
| P04104 | Keratin, type II cytoskeletal 1 (GN=KRT1) | 8.9 |
| P02535 | Keratin, type I cytoskeletal 10 (GN=KRT10) | 3.5 |
| P11276 | Fibronectin (GN=FN1)* | 0.04* |

**Fraction 12 of SV gradient from C2C12 in Triton X100 Na-Deoxycholate (Figure 3)**
| Accession | Description | Percentage of Contamination |
| --- | --- | --- |
| P11087 | Collagen alpha-1(I) chain (GN=COL1A1) | 55.4 |
| P04104 | Keratin, type II cytoskeletal 1 (GN=KRT1) | 14.7 |
| Q01149 | Collagen alpha-2(I) chain (GN=COL1A2) | 8.1 |
| Q61554 | Fibrillin-1 (GN=FBN1) | 7.4 |
| P11276 | Fibronectin (GN=FN1) | 4.7 |
| P02535 | Keratin, type I cytoskeletal 10 (GN=KRT10) | 4.3 |
| Q05793 | Basement membrane-specific heparan sulfate proteoglycan core protein (GN=HSPG2) | 2.0 |
| P62806 | Histone H4 (GN=HIST1H4A) | 1.6 |
| Q02788 | Collagen alpha-2(VI) chain (GN=COL6A2) | 1.1 |
| P04247 | Myoglobin (GN=MB) | 0.7 |

\* Extracted Ion Chromatogram not calculated as protein identification occurred below limit of detection (LOD). Arbitrary LOD threshold value applied for graphical representation.

Electron microscopy of fractions containing highly pure PrP^res^ (fractions #13 for sarkosyl and #11 for TD) showed typical prion rod structures when sarkosyl was used but more blob-like structures when TD was used (Fig 5B, C). In PTA-precipitated samples, these prion rods structures were present but mostly intermingled with other fibrillar structures with the characteristic striated ultrastructure of collagen (Fig 5D). Unfortunately, FN is undetectable by conventional negative staining [27].

### 4. FN negatively correlates with PrP^Sc^ in dC2C12 cultures

The distribution of FN, the most abundant contaminating protein as suggested by immunoblot, was analyzed by ICC in dC2C12 cultures. At day 20 post-differentiation in NBH-treated dC2C12 cultures, the ICC showed FN staining as a filamentous mesh-like pattern (Fig 6A). The FN staining was intense in areas with an abundance of reserve cells, while areas with a predominance of myotubes had no or very faint FN staining, likely just background from lower layers. The same FN pattern was observed in RML-infected dC2C12 cultures. Interestingly, an inverse correlation between PrP^Sc^ and FN staining was observed. Areas with intense FN staining were completely devoid of PrP^Sc^ and, conversely, areas with PrP^Sc^ showed very low or no FN staining. Based on cell morphology, the areas with low FN staining and intense PrP^Sc^ punctate were associated with myotubes, while intense fibrillary patterns of FN and an absence of PrP^Sc^ co-localized with reserve cells (Fig 6B). The amount of FN decreased from day 7 to 14 post-differentiation (Fig 7A) indicating changes in FN expression during the maturation of the culture. Immunoblotting analysis showed no significant differences in the levels of FN between NBH and RML-infected dC2C12 cultures at day 20 (Fig 7B), suggesting that RML infection did not affect FN metabolism.

**Fig 6.**
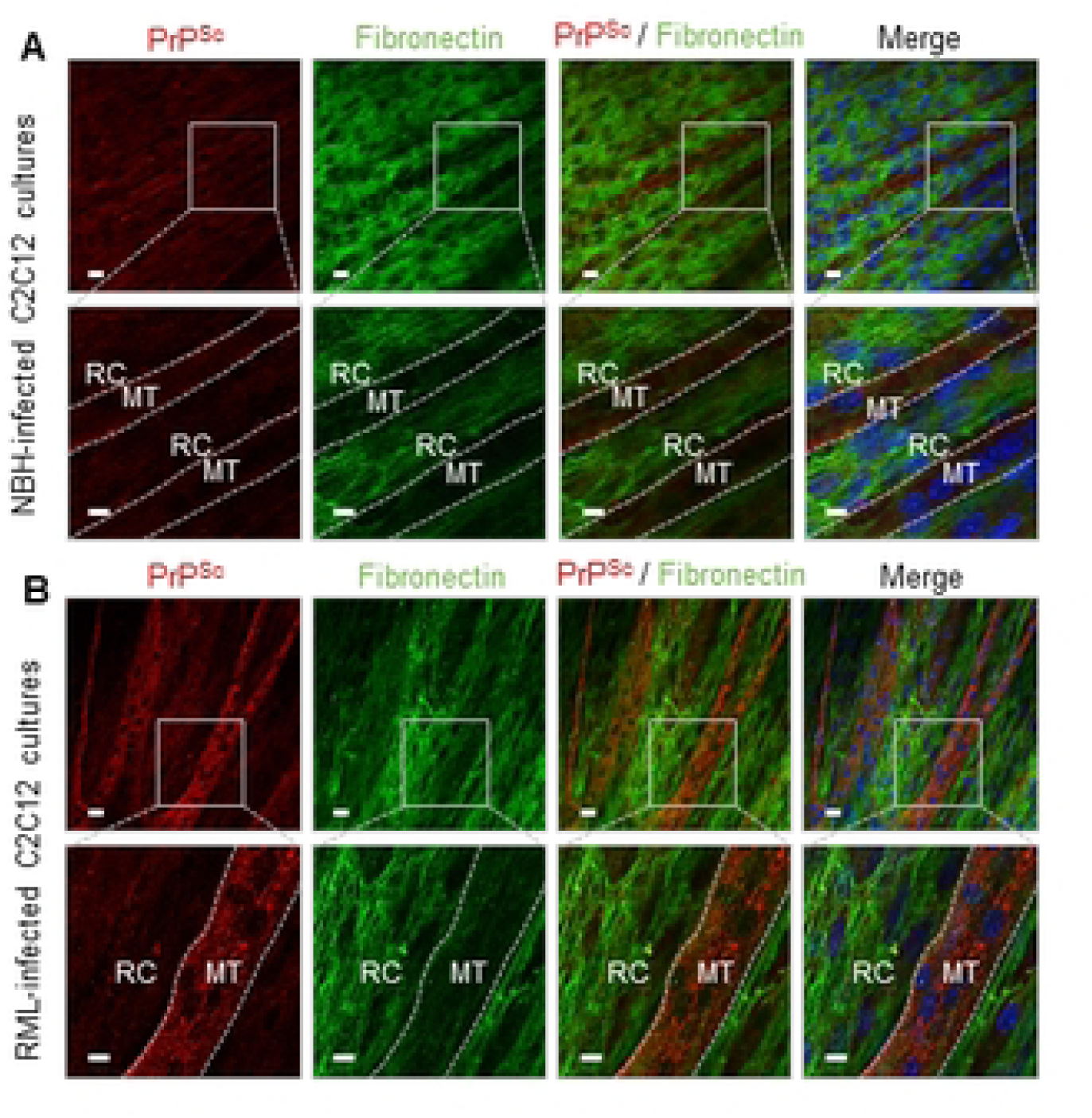
Detection of PrP and FN proteins in muscle-derived cells. (A) Immunocytochemical detection of PrP^Sc^ (red) and FN (green) in NBH-treated and RML-infected dC2C12 cultures at day 20 p.i. No co-localization of the two proteins was observed in either the NBH-treated or RML-infected cultures. Nuclei in blue. MT, myotube; RC, reserve cells. Scale bar, 20 µm and 40 µm in the boxed images.

**Fig 7.**
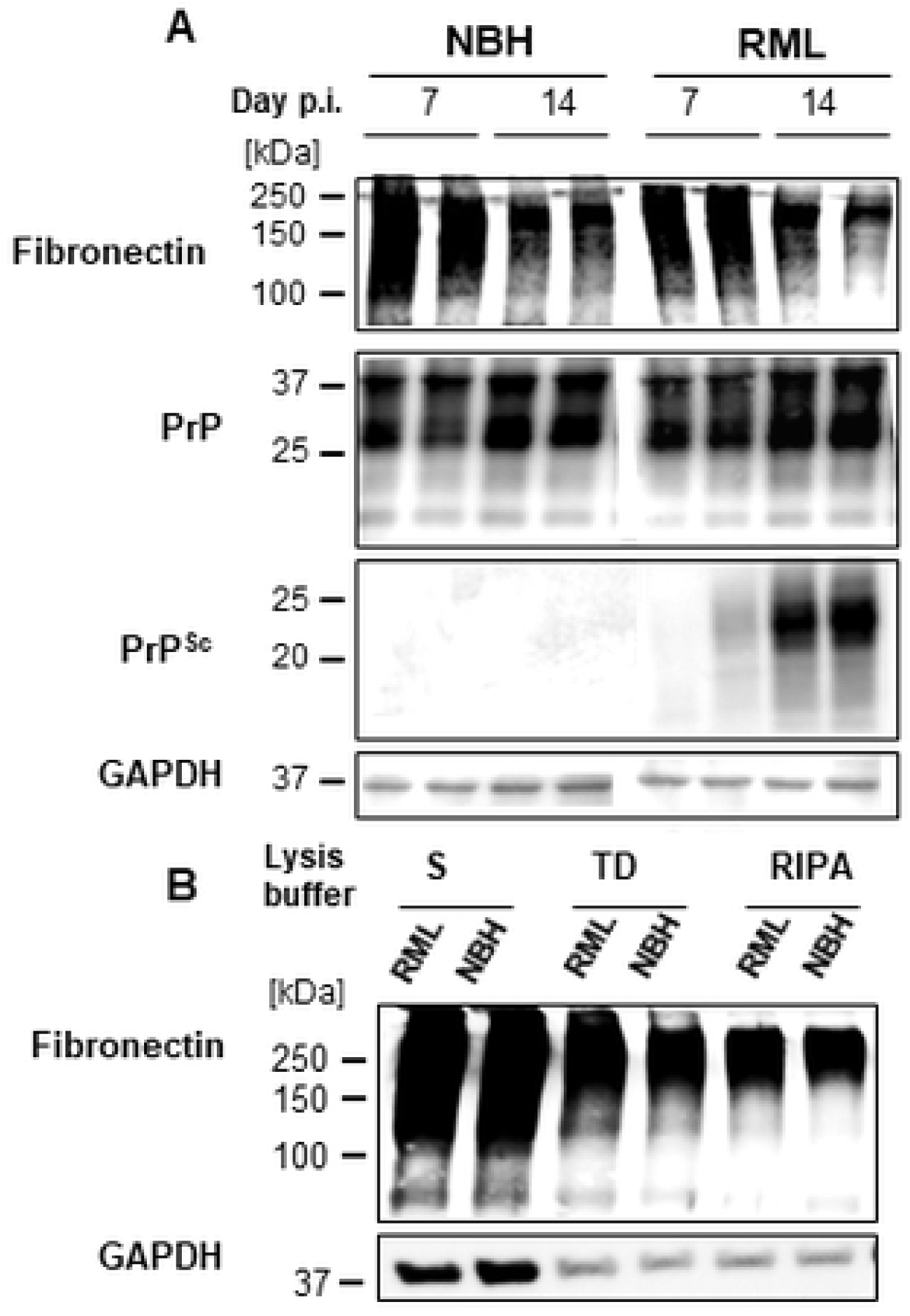
Time course of PrP^res^ and FN in dC2C12 cultures. Immunoblot analysis of PrP^res^ and FN at 7 and 14 days p.i. show the negative correlation of the presence of these proteins with time. (A) FN decreases from day 7 to 14 in both NBH-treated and RML-infected cultures whereas PrP^res^ increases with time. (B) No differences in FN levels between NBH and RML-infected cultures at day 20 are detected. The detergents used in the homogenization of the lysates yield different amount of FN by WB, sarkosyl yielding the larger FN signal.

### 5. FN does not interact with PrP^C^

An interaction between FN and PrP^C^ may hide the substrate for PrP^Sc^ replication explaining the resistance of reserve cells to prion infection. We used chemical cross-linking to study this interaction. After treating NBH-infected dC2C12 cultures with 2% formaldehyde, the samples were analyzed by immunoblotting. Signal for PrP^C^ was detected at the size corresponding to PrP monomers but also at 150-250 kDa (S3 Fig). These HMW bands disappeared with temperature cross-linking reversion. The intensity of the signal for monomeric PrP^C^ after reversion was almost the same as the addition of the signals for monomeric and complexed PrP^C^ before boiling of the sample. FN band intensity was the same after cross-linking and at different times of reversion suggesting no cross-linking of FN with other proteins under our experimental conditions. The primary interactor of PrP^C^ at day 10 post-differentiation was neural cell adhesion molecule (NCAM) (S3 Fig) as previously described [28]. Immunoblot of cross-linked samples with anti-NCAM antibody showed bands at the same positions as PrP^C^, between 150 and 250 kDa. These bands also disappeared by boiling the sample for 5, 15 and 30 minutes (S3 Fig). Finally, we evaluated the *in vitro* interaction of recombinant PrP (recPrP) with different aggregation states of purified FN by surface plasmon resonance. FN aggregates were generated under various conditions of agitation and temperature, and the aggregation state was determined by dynamic light scattering. None of the FN preparations (monomeric, 240kDa; dimeric, 420 kDa; tetrameric, 714 kDa; and fibrillar) showed binding to recPrP.

### 6. Dexamethasone treatment renders reserve cells permissive to prion infection

To confirm that the rich FN matrix present in reserve cells is responsible for their refractory status to prion infection, dC2C12 cultures were exposed to dexamethasone (Dex). This corticosteroid induces atrophy in myotubes [29–32] by decreasing protein synthesis and increasing the rate of protein catabolism [29,31,33] rendering a culture enriched in reserve cells (Fig 8A). Transcription factors regulating myogenic differentiation were examined by immunoblotting analysis. The expression of MyoD (a myogenic regulatory factor) and Pax7 (a paired box transcription factor) were decreased by Dex treatment regardless of prion infection (Fig 9A). This myogenic inhibition by Dex supports the immunocytochemical observation of myotube atrophy in the differentiated C2C12 culture shown in Fig 8.

**Fig 8.**
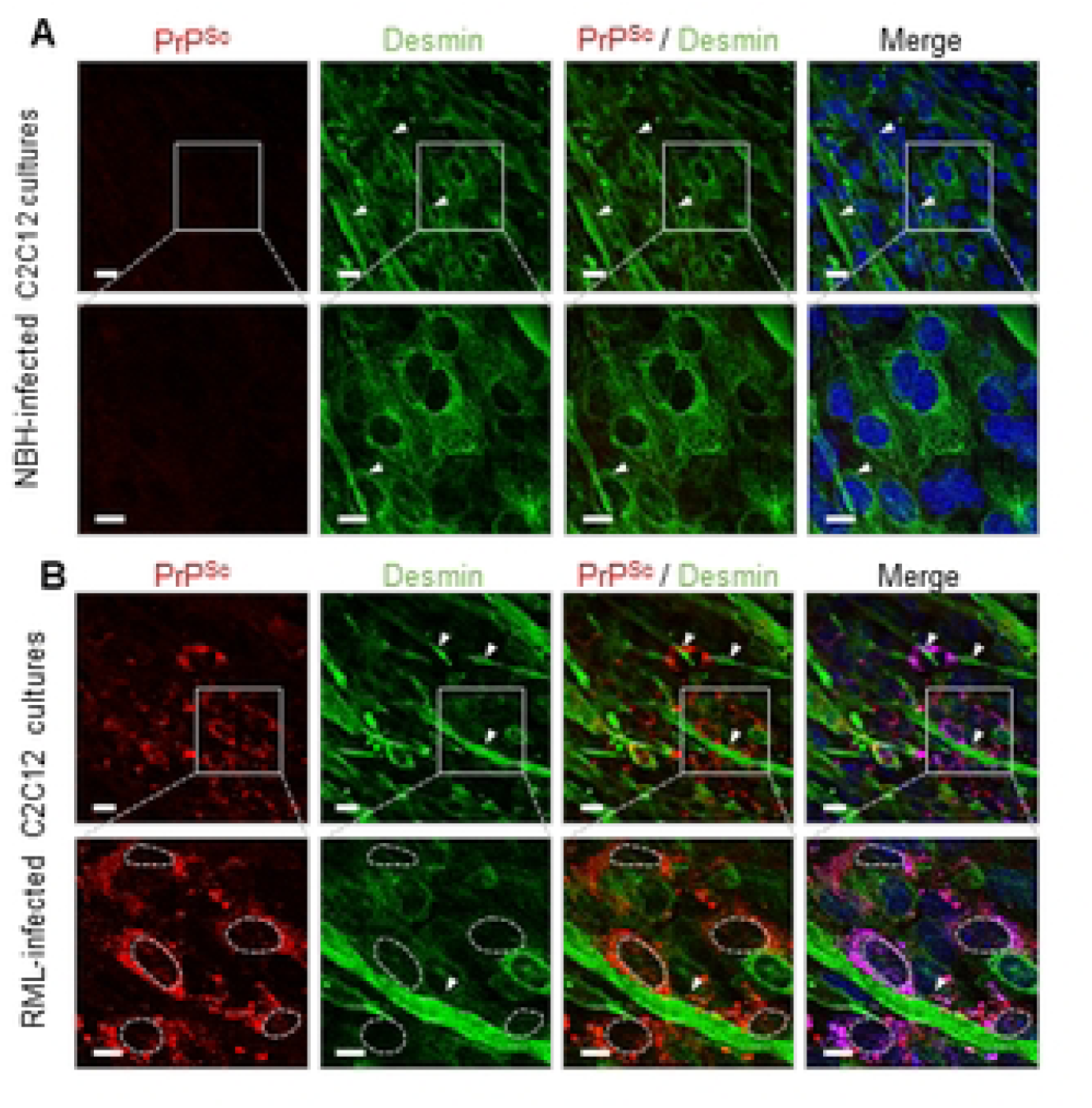
Dexamethasone effect on dC2C12 cultures confirms the inverse correlation between prion accumulation and FN. Differentiated C2C12 cultures were treated with Dex (10 µM) at day 7 post-exposure to NBH or RML prions. (A) Immunocytochemical detection in cultures at 21 days p.i. of PrP^Sc^ (in red) and desmin (in green) reveal some atrophic myotubes remaining caused by Dex treatment and an abundance of mononucleated cells (indicated by arrowheads). PrP^Sc^ was observed, for the first time, associated with the mononucleated cells as gross particulates. Scale bar, 10 μm.

**Fig 9.**
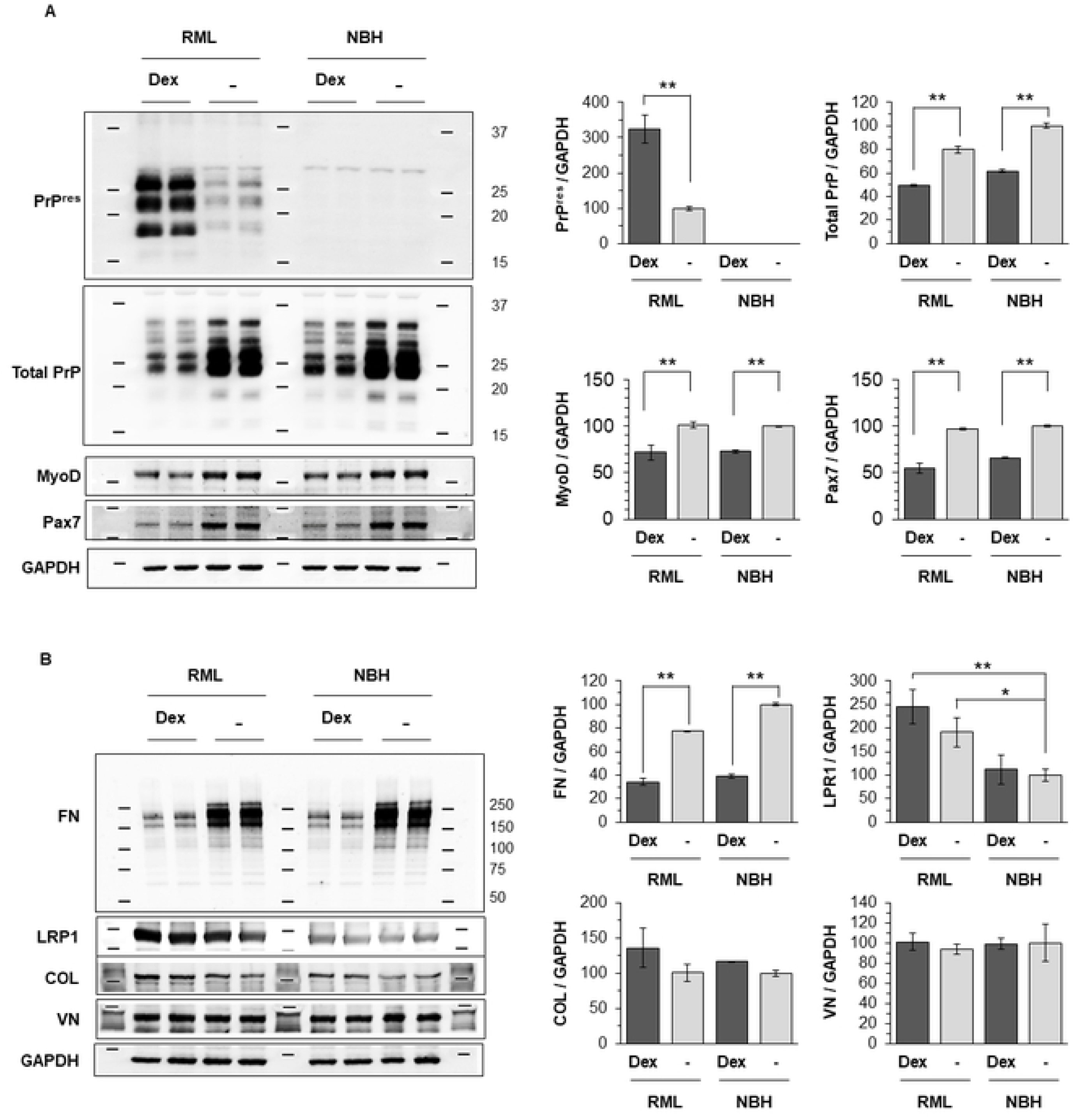
Protein expression of PrP and ECM components in dC2C12 at 20 days p.i. treated with Dexamethasone. dC2C12 culture was exposed to RML prions or NBH, and then, at 7 days post-infection, treated with Dex (10 μM in ethanol) or mock control (0.25% ethanol). The cells were harvested at 20 days post-exposure and analyzed by immunoblotting. Dexamethasone treatment of NBH and RML infected dC2C12 reduces (A) total PrP levels and (B) FN with a concomitant increase in (A) PK-resistant PrP compared with non Dex-exposed cultures. (A) The expression of MyoD and Pax7, myogenic regulatory factors, was reduced while (B) other extracellular matrix (ECM) proteins such as collagen I and vitronectin did not change. Conversely, LRP1, reported to mediate endocytosis of PrP^C^ in neuronal cells, was increased proportionally to the accumulation of PrP^res^. The immunoblot intensity was normalized by GAPDH loading control. Error bars represent SD. **, P<0.01 in comparison with mock treatment control. Error bars represent SD. *, P<0.05 and **, P<0.01. FN, fibronectin; protein 1; LRP1, low-density lipoprotein receptor-related protein 1; COL-I, collagen I; VN, vitronectin; GAPDH, Glyceraldehyde 3-phosphate dehydrogenase.

Upon infection, Dex-treated cultures showed PrP^Sc^ in relation to mononucleated cells with morphologies resembling reserve cells. The pattern of PrP staining changed from fine-punctate or diffuse in non-Dex cultures to thick intra-cytoplasmatic granules or coalescing aggregates surrounding the nucleus in Dex-exposed cultures (Fig 8). Some shrunken myotubes that also displayed PrP^Sc^ were still present. Immunoblotting analysis revealed an increase in total PrP^Sc^ signal concomitant with a decrease in the FN levels in Dex-RML treated cultures (Fig 9A); the decrease in FN also appeared in Dex-NBH cultures and was significant at two different time points (10 and 20 days p.i.; Fig 9B). Unlike FN, there were no changes in other ECM proteins like collagen and vitronectin. The increased PrP^Sc^ signal after Dex treatment, even at the two different time-points, was not correlated with any increase in the total PrP. In fact, the total PrP levels decreased (Fig 9B).

### 7. Looking for PrP^Sc^ and FN in tissues from naturally infected sheep

We then investigated if the inverse correlation between FN and PrP^Sc^ deposits observed *in vitro* was also present *in vivo*. In a previous study, we examined the extent, location and pattern of PrP^Sc^ deposition in several tissues from naturally infected scrapie sheep [10]. We further analyzed those samples to determine the distribution of FN in relation to PrP^Sc^ deposition, specifically in muscle tissues and in the targeted tissues of major prion diseases: CNS, enteric nervous system (ENS) and adrenal gland. PrP^Sc^ and FN immunodetection was performed on serial sections of tissues by IHC.

Three different types of muscles were examined: skeletal, cardiac striated and smooth muscle. In skeletal muscle, FN immunostaining was detected at the endomysium as a fine punctuate stain that sheathed each individual myofiber; no PrP^Sc^ was found associated with myofibers (Fig 10A, B) as previously described [9]. The presence of FN at the endomysium also appeared in smooth muscles from the digestive tract. The spindle-shaped cells from the muscularis propria of the digestive wall were enclosed by a FN-positive sheath of endomysium and were negative for PrP^Sc^ (Fig 10C, D). PrP^Sc^ in these samples was associated with the enteric nervous system, largely at the myenteric plexus, between the longitudinal and circular layers of the muscularis, consisting of ganglia interconnected by bundles of nerve fibers. In these ganglia, neural and glial elements were densely packed with scarce extracellular space that was negative for FN (Fig 10E, F). However, as previously reported [34], the perineurium surrounding each PrP^Sc^ positive ganglion was positive for FN (Fig 10E, asterisk), which would constitute a physical barrier from the adjacent smooth muscle layers, maintaining the negative correlation of FN and PrP^Sc^ in the gastrointestinal wall. In contrast to the skeletal and smooth muscle, no immunostaining for FN was observed in the myocardium, suggesting that the protein composition of the cardiac endomysium differs from that of other types of muscle (see S4 A Fig). Furthermore, IHC analysis revealed a lack of PrP^Sc^ in this tissue (see S4 B Fig).

**Fig 10.**
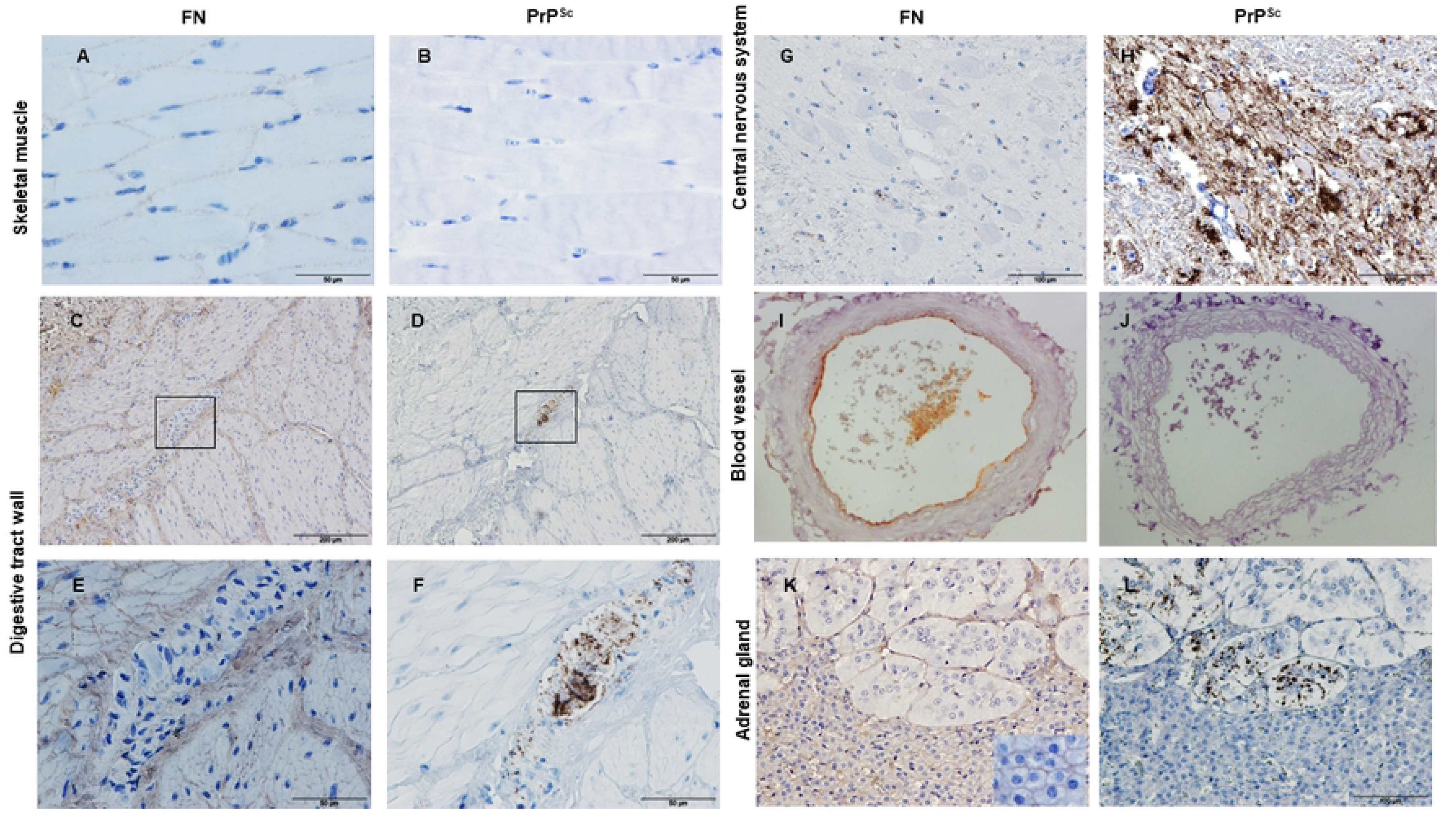
PrP^Sc^ is inversely correlated with FN in tissues from naturally scrapie-infected sheep. (A, C, E, G, I, K) Immunohistochemical detection of FN and (B, D, F, H, J, L) PrP^Sc^ in serial sections of tissues from naturally scrapie-infected sheep. (A, B) In striated skeletal muscle, fine punctae of FN inmunolabelling are observed at the endomysium sheath of the myofibers. (C, D, E, F) In smooth muscle layers of the digestive tract, FN is found at basal lamina of each myocyte with PrP^Sc^ restricted to the myenteric plexus, surrounded by a FN positive perineurium (asterisk). (G, H) The central nervous system shows intense PrP^Sc^ accumulation. (I, J) Blood vessel displayed FN at the basal lamina of endothelium and at the adventitia. (K) The adrenal gland shows FN staining with a diffuse pattern in the entire adrenal cortex at intercellular space (inset: detailed FN intercellular staining); (L) PrP^Sc^ accumulation is associated with chromaffin cells at the adrenal medulla.

As expected, no FN was detected in the CNS tissue sections examined (Fig 10G, H) [35]. However, the lumen of the vessels was positive for FN staining due to the antibody recognizing indistinctly soluble (plasma) and insoluble (cellular) FN. This staining of soluble FN appeared in vessels for all the tissues examined. In addition, transverse sections of the vessels had clear FN staining as circular patterns in the connective tissue of tunica intima (subendothelial) and tunica adventitia (Fig 10I, J).

Apart from the central and peripheral nervous tissues, the adrenal gland was also examined as it is positive for PrP^Sc^ in all major prion disorders (scrapie, BSE, CWD, CJD) [36–39]. FN staining was found in all intercellular spaces of the cells in the adrenal cortex whereas in the medulla, chromaffin cells lacked any intercellular FN staining. These cells were scattered in clusters and FN staining was present around the clusters, but not within (Fig 10K). Again, there was an inverse correlation between FN and PrP^Sc^. In the adrenal cortex, the highest intercellular FN immunostaining was detected with no PrP^Sc^ deposits. PrP^Sc^ was restricted to within the clusters of chromaffin cells either at nerve fibers/endings or as intracytoplasmic deposits in chromaffin cells [40] where no FN was present (Fig 10L).

In scrapie, the lymphoid tissue is of particular interest, while in other prion disorders like BSE, this tissue is not or minimally involved in prion pathogenesis. In general, mild diffuse FN labelling was observed in the medulla of the lymph nodes, none coexisting with PrP^Sc^ that was instead located in primary/secondary follicles of the lymph node cortex (S4 C, D Fig).

## Discussion

### Prion tropism in reserve cells and myotubes

The accumulation of PrP^Sc^ in striated muscle tissue of prion-infected animals is associated with nerve structures rather than myofibers [10,12–15], suggesting that there is an inefficient environment for prion replication and/or accumulation in this cell type. Conversely, myotubes, the myofiber counterpart in dC2C12 cell culture, become susceptible to prion infection when they mature from their prion-resistant myoblast precursors. In addition to myotubes, dC2C12 culture also contain reserve cells that remain undifferentiated and prion resistant [21,22,41]. Here we explored the cellular factors that account for the different prion susceptibility of these two muscle cell types.

We identified FN, fibrillin and collagen as co-purifying proteins with PrP^Sc^ from dC2C12 culture. The consistent co-precipitation and co-migration of these structural proteins with PrP^Sc^ under a wide range of experimental conditions strongly suggests their interaction. In addition, these co-purifying proteins reduced PrP^Sc^ infectivity in SSCA. ICC analysis of infected dC2C12 cultures showed an inverse correlation between FN and PrP^Sc^ deposition, with high levels of FN at ECM surrounding prion-resistant reserve cells and low levels surrounding prion-susceptible myotubes. This supports the hypothesis that these proteins interfere with prion replication.

Interestingly, induction of myotube atrophy and reduction of FN levels by treatment with Dex made reserve cells permissive to prion infection. This further suggests that these structural proteins (normally found in systemic ECM), especially FN, protect from prion infection through direct or indirect interaction with PrP^Sc^. The atrophic status of myotubes treated with Dex was also confirmed by expression levels of transcription factors regulating myogenic differentiation, MyoD and Pax7. MyoD, a skeletal muscle-specific transcription factor, is crucial for the balance between proliferation and withdrawal from the cell cycle, resulting in terminal differentiation or quiescence [20,42,43]. MyoD-positive cells align and fuse to form multi-nucleated myotubes, while MyoD-negative cells remain undifferentiated and mononucleated (reserve cells) [44]. Pax7, a paired box transcription factor, is a marker for reserve cells in the activated and proliferative states and plays a role in the maintenance of reserve cell pool [45]. A cross-inhibitory interaction between MyoD and Pax7 affect cell fate determination [46]. Since activated reserve cells, which are proliferating and differentiating, co-express MyoD with Pax7 [45,47,48], the reduction in both MyoD and Pax7 detected in the differentiated C2C12 culture (Fig 9A) might indicate that myotube atrophy was initiated partially by inactivation of reserve cells from the myogenic regeneration processes.

Several studies have indicated that myoblasts, which are resistant to prion infection [22,41], synthesize and organize FN in a rich surface-associated matrix that is lost in myotubes during differentiation [49,50]. In dC2C12 cultures, the myotube formation requires a switching of the cell-ECM interaction that involves the degradation of 90% of FN by membrane-type 1 matrix metalloproteinase (MT1-MMP) during the elongation phase [47]. Our ICC results show that in dC2C12 culture, reserve cells, like myoblasts, display this rich FN surface-associated matrix whereas myotubes do not.

GPI-anchored proteins, such as PrP^C^, can perform signaling roles integrating the ECM with the cytoskeleton. For instance, binding of PrP^C^ to extracellular matrix protein laminin was reported to induce neurite maintenance and neuronal differentiation [51,52] and high affinity interaction with vitronectin supports axonal growth [53]. An interaction of FN with PrP^C^ could result in a protective role by sequestering the substrate for PrP^Sc^ replication, similar to the inhibitory effect on prion replication *in vivo* caused by anti-PrP monoclonal antibodies [54]. As previously reported, we observed no evidence of a direct FN-PrP^C^ interaction [53].

Our results provide additional evidence that ECM proteins play a crucial role in cellular permissibility to prion infection. Transcriptomic analysis of prion-resistant revertant cells in comparison with their susceptible counterparts identified a gene network for ECM components affecting prion replication *in vitro* [55,56]. In particular, a rich FN ECM was characteristic of the inhibitory phenotypic state for prion replication in N2a cell culture [56]. Susceptible N2a subclones had low FN expression levels whereas resistant subclones overexpressed this protein. As we observed in dC2C12 culture, confocal images of the susceptible N2a culture showed an inverse correlation between FN and PrP^Sc^, and silencing FN expression augmented the rate of prion replication [56,57].

### Fibronectin and prion accumulation in muscle

It is noteworthy that, in contrast to *in vitro* myotubes that replicate prions very efficiently [41], myofibers, their natural counterparts, do not appear to replicate prions in muscle tissue of natural disease. Interestingly, intramuscular inoculations of prions in transgenic mice corroborates the capability of myofibers to produce high titers of nascent prions [58] suggesting that extracellular factors play a significant role in maintaining the resistant status of myofibers *in vivo*. In the context of our findings, this prion resistance observed *in vivo* could be related to the presence of a thin pericellular layer of FN in the ECM that surrounds myofibers in muscle (endomysium) acting as a physical barrier against PrP^Sc^, that is absent in dC2C12 myotubes [59,60]. Different locations of nuclei in myotubes and myofibers may account for the differences in FN content between these two types of cells. The deposition of extracellular FN at the periphery of myofibers triggers the peripheral positioning of their nuclei, in close contact with the plasma membrane [61], whereas the central position of the nuclei in myotubes may be a consequence of the lack of pericellular FN. In addition to the pericellular FN matrix, there are numerous structures of connective tissue interposed between the entry points of prions to muscle, the neuromuscular spindles and nerve fascicles [9,10,62] and the myofibers. Specifically, myofibers are mantled by up to three layers of ECM: endomysium, perimysium and epimysium. Therefore, prions from nerve fascicles need cross at least two of the layers, perimysium and endomysium, to reach the cells, bearing in mind that the nervous structures are confined by their own connective tissue, endoneurium and perineurium. The penetration of prions to myofibers *in vivo* would then be hindered by a large extension of connective tissue and foremost the FN matrix at the endomysium that is interposed between the prion entry points to the tissue and the myofibers.

We found an inverse correlation between FN and PrP^Sc^ accumulation in the three types of muscle examined as well as in other tissues from a natural model of prion disease, ovine scrapie. In agreement with the literature [61,63], we confirmed the presence of a robust FN immunostaining in skeletal and smooth muscle but not in cardiac muscle. In the case of cardiac muscle, the literature describes the presence of low levels of FN at the interstitium of the healthy heart [64–66] indicating that collagen and elastic fibers are the more abundant fibers. Interestingly, in a previous study [10], we rigorously examined for PrP^Sc^ in cardiac and skeletal muscles, detecting an unexpected proportion of hearts that were positive by ELISA (4/16), whereas no skeletal striated muscles (0/16) were positive by this technique. IHC examination of those hearts showed, for the first time a few dots of PrP^Sc^ at intercellular spaces. In addition to our results from scrapie infected animals, Jewell JE; *et al.* 2006 reported the detection of PrP^Sc^ in cardiac muscle of two species of cervid with CWD (7/18) describing linear patterns along the edge of muscle cells[67]. Morphologically, the cardiomyocytes are more similar to dC2C12 myotubes than skeletal myofibers. Cardiomyocytes, which have central nuclei like myotubes, might be devoid of a rich FN endomysium, thus explaining the PrP^Sc^ positive hearts.

Like skeletal muscle, smooth muscle cells remain negative for PrP^Sc^ despite their close proximity to PrP^Sc^ positive nervous structures. Among the two layers of smooth muscle in the gastrointestinal wall, run the myenteric and submucosal plexi, which show abundant PrP^Sc^ accumulation. The intestinal plexi belong to enteric nervous system (ENS), through which the prions transit and reach the CNS after oral exposure. Thus, we hypothesize that, along the digestive tract, the smooth muscle layers of the wall remained negative because the FN at basal lamina of each individual myocyte and at perineurium surrounding the plexi may provide a barrier that prevents the spreading of prions between layers.

### Fibronectin and prion accumulation in nervous system and other peripheral tissues

The CNS harbors the vast majority of prion infectivity, lesions and PrP^Sc^ accumulation in all prion disorders. As expected, our IHC examination of the CNS from scrapie-infected animals confirmed high levels of PrP^Sc^ and undetectable levels of FN. The ECM in brain has a different composition than the systemic ECM [68], being richer in proteoglycans with minimal amounts of FN and collagen [69–71]. The peripheral nervous system (PNS), the primary tissue for neuroinvasion in acquired prion disorders, typically lacks FN in mature stages, except during injury or regeneration [72]. Interestingly, in both the CNS and PNS, FN is the fibrillar protein present at developing stages and replaced by laminin at maturation, these changes in ECM composition [72–74] would favor prion replication.

In vCJD, CWD and scrapie, prion replication in lymphoid organs such as gut-associated lymphoid tissue (GALT), lymph nodes and spleen, is necessary for effective neuroinvasion [75–78]. The lymphoid tissue examined in our study, whether lymph nodes or mucosa-associated lymphoid tissue, revealed PrP^Sc^ positive lymphoid follicles primarily at the light zone of the germinal center, whereas the mantle and interfollicular areas, expressing more FN, were devoid of PrP^Sc^.

Apart from nervous and lymphoid tissue, the adrenal gland is the only other tissue that exhibits PrP^Sc^ deposition in all acquired well-studied prion disorders: scrapie, BSE, CWD, vCJD. [36,39,40,79]. Despite being an endocrine organ, the adrenal medulla is considered a modified neural tissue functioning like a sympathetic ganglion, in fact, medulla and cortex of the adrenal gland differ in their embryonic origin, with a neuroectodermal and mesodermal origin respectively. The different tissue composition of these regions within the organ might explain the distribution of these two proteins, with PrP^Sc^ accumulated in the medulla, and FN, largely concentrated to the cortex. The selective prion presence in the medulla would respond to the FN lacking featured of this nervous region.

Spread of prions through the peripheral nervous system (PNS) is well recognized, but the hematogenous route may be a parallel or alternative pathway that contributes to the peripheral distribution. However, despite the presence of considerable amounts of infectivity in blood, there is limited contribution of this route to distribution of PrP^Sc^. We consistently observed linear FN immunostaining surrounding the entire diameter of blood vessels, corresponding to the basal lamina of the endothelia. This may explain the low contribution of the hematogenous route to prion spread compared to dissemination via the peripheral nerves [80].

We believe that the reduction of the glycoprotein FN in the ECM during the developing nervous system and in C2C12 cultures favors prion replication. In the systemic ECM, FN polymerizes to higher order fibrils that assemble into an insoluble sticky matrix, limiting prion replication and /or transport. Although other factors may influence prion tropism and spread [81], our results strongly suggest that a rich FN extracellular matrix protects cells from prion infection not only by acting as a physical barrier but also as a biochemical barrier, interacting with PrP^Sc^ and inhibiting its uptake into cells. If we consider that the contact between infectious PrP^Sc^ particles and normal PrP^C^ takes place at the cell surface, an ECM that does not favor this interaction would be the first barrier against prion replication. These findings provide new insights into prion tropism in different tissues.

## Material and methods

### Infection of Cell cultures and sample preparation

C2C12 (CRL-1722) myoblast cells were purchased from the American Type Culture Collection and maintained as previously described by Herbst *et al*., 2013 ([19]). Following expansion of the myoblast cells, aliquots stored in liquid nitrogen and further expanded as needed. Myoblasts were grown in Dulbecco’s Modified Eagle Medium, 10% fetal bovine serum with penicillin and streptomycin (PS). Myoblasts were differentiated when confluent by switching to differentiation medium; DMEM, 10% horse serum (HS), and PS. Four days after myotubes first appeared, infections were initiated by the addition of prion-infected brain or normal brain homogenates (hereafter named NBH-treated) diluted in differentiation media (DMEM, 10% HS, 1% PS) at final concentration of 0.01%. Media was changed one day after prion infection to remove residual inoculum and every two days subsequently. Myotube atrophy was induced by Dex (InvivoGen, CA, US, tlrl-dex) treatment at 100 nM concentration 7 days after prion infection. Cells were harvested 20 days post-infection in RIPA buffer or cell lysis buffer containing 1mM PMSF. Different cell lysis buffers were used to examine their efficacy in PrP^Sc^ extraction containing 50 mM Tris, 150 mM NaCl, 1mM EDTA but with different detergents: Sarkosyl from 0.5 to 5%, Triton X-100 from 0.5 to 1%, Na-Deoxycholate 0.25 to 0.5%). Some cells were harvested in PBS Gibco and physically disrupted by sonication (1 minutes of sonication and cooling on ice 1 minute for three times). Lysis buffer used: *RIPA Lysis buffer:* 50 mM Tris base, 150mM NaCl, 1mM EDTA, 1% CA-360, 0.25% Na-deoxycholate. *Lysis buffer containing sarkosyl*: 50 mM Tris base, 150mM NaCl, 1mM EDTA, 1% CA-360, 2% sarkosyl. *Lysis buffer containing 1% TRITON X100*: 50 mM Tris base, 150mM NaCl, 1mM EDTA, 1% CA-360, 1% Triton X100. *Lysis buffer containing TD0.5%:* 50 mM Tris base, 150mM NaCl, 1mM EDTA, 1% CA-360, 0.5% Triton, 0.5% Na-deoxycholate.

#### Prion inoculum preparation

C57BL/6 mice were intracerebrally inoculated with RML mouse prions. The animals were euthanized at the clinical stage of the disease. Brains from clinically affected mice were homogenized in phosphate buffered saline (PBS) and centrifuged at 100 g for 5 min to remove cellular debris. Total protein concentration was adjusted to 2 mg/mL based on BCA protein assay (Pierce, MA, US, 23235) and samples were stored at -80°C until use. Animal inoculation studies were performed at the University of Alberta following the Canadian Council on Animal Care guidelines, reviewed by the ACUC-HS, protocol #914)

### Immunocytochemistry

Cells cultured on SlideFlasks (Nunc, NY, US, 170920) were fixed in paraformaldehyde (4%, pH 7.4) for 15 min and permeabilized with PBS containing Triton X-100 (0.1%). To expose epitopes of PrP^Sc^, cells were treated with guanidine thiocyanate (3 M GdnSCN) for 2 hours at 4°C. The fixed cells were blocked with 1% bovine serum albumin (BSA) in PBST (PBS with 0.1% Tween 20) for 30 min. PrP^Sc^ was detected with anti-PrP mAb, SAF83 (Cayman, MI, US, 189765) in GdnSCN-treated cells. Desmin, and fibronectin (FN) were detected with anti-desmin pAb (Abcam, Cambridge, UK, ab15200) and anti-FN pAb (Abcam, ab2413), respectively. To visualize the target molecules, cells were then incubated with appropriate fluorescent-conjugated (Alexa Fluor 488 or Alexa Fluor 594, Invitrogen) secondary antibodies. Nuclear counterstain was performed with 4’,6-diamidino-2-phenylindole (DAPI) fluorescent stain (Invitrogen, P36935). Cells were analyzed by confocal microscopy (LSM700 laser scanning microscope, Zeiss, Jena, Germany) using Z-stack functions with identical imaging settings.

#### Measurements of myotube width

38 myotubes from micrographs at 40x magnification were examined per culture and the width value for each myotubes was calculated as the average of 10 measurements per myosegement; Chi squared equals 22.021 with 4 degrees of freedom. The two-tailed P value equals 0.0002.

### PTA based purification protocol

Cell lysates were homogenized by numerous pipetting and digested with Benzonase-nuclease (100 U/ml; Sigma-Aldrich) before clarification at 500xg for 10 minutes. The aliquots were frozen at -80°C until processed. A purification protocol, mainly based on proteinase digestion followed by precipitation with sodium phosphotungstic acid (NaPTA) in the presence of Sarkosyl, was performed. The use of NaPTA was adapted from Safar and colleagues [24] as described previously [36]. Briefly, thawed aliquots of cell lysates harvested in different lysis buffers were incubated with 2% Sarkosyl for 1 hour, and subsequently digested with proteinase K or pronase E (100ug/ml) for 30 minutes. After stopping the reaction with 2mM PMSF and 10mM EDTA, the samples were incubated with 0.3% NaPTA (pH 7.0) for 1 hour and centrifuged at 16,000xg at 4°C for 30 minutes. The pellet was resuspended in 2% sarkosyl and incubated overnight. Samples were adjusted to 0.3% (w/v) NaPTA and incubated for 1 hour, obtaining the final pellet by centrifugation at 1800xg at 4°C for 30 minutes. All digestions and incubation were performed at 37°C with vigorous agitation.

### Velocity sedimentation

The sample fractionation was performed by iodixanol (OptiPrepTM) gradients applying sedimentation velocity (SV) techniques as previously described by Tixador *et al.,* [26]. Optiprep density gradient medium (Sigma) was dissolved in a buffer containing 25 mM HEPES pH 7.4, 150 mM NaCl, 2 mM EDTA, 1 mM DTT, 0.5% Sarkosyl to perform the different concentrations of iodixanol, after manually layering, the gradients were allowed to stand overnight at 4°C to produce the continuous gradient. Gradient linearity was verified by refractometry. Cell lysates or brain homogenates (from 200 to 500ul and 100ul, respectively) were layered on a 3.6ml continuous 12.5-25% iodixanol gradient. The ultracentrifugations were carried out in a swinging-bucket SW-55 rotor using an Optima LE-80K ultracentrifuge (Beckman Coulter), at 285000xg 45 minutes. Finally, the gradients were manually segregated into 14 equal fractions, numbered from top to the bottom, and aliquoted for SDS page, SSCA and EM analysis.

The purified samples and fractions were separated by reducing by SDS-PAGE followed by silver staining or by blotting on PVDF membranes (BioRad) and blocked with 5% BSA in Tris-buffer saline supplemented with 0.05% Tween20 (TBST). The blot was incubated with primary (overnight at 4C) and secondary antibodies (1 hour) in 5% BSA/TBST. Detection was performed with Clarity™ Western ECL Blotting Substrate (BioRad). The antibodies used were: anti-PrP mAb, SAF83 (Cayman, 189765); anti-MyoD mAb (ThermoFisher, IL, US, MA1-41017); anti-collagen type I alpha1 chain (COL1A1) pAb (Invitrogen, PA5-29569); anti-vitronectin (VN) pAb (Santa Cruz, sc-28929); anti-glyceraldehyde 3-phosphate dehydrogenase (GAPDH) mAb (Abcam, ab9484); anti-desmin pAb (Abcam, Cambridge, UK, ab15200) and anti-FN pAb (Abcam, ab2413).

### Statistical Analysis

Statistical analysis for immunoblot results was performed using the independent sample t test and one-way analysis of variance (ANOVA) followed by post hoc with the least significant difference (LSD) analysis. Statistical analysis of all data was performed using IBM SPSS Statistics version 24 software.

### Negative stain electron microscopy

Transmission electron microscopy was performed using a TF20: Tecnai G2 200kV TEM (FEI) operating at 200 kV. Five μL of sample (no dilution) were adsorbed on freshly glow-discharged 200-mesh formvar/carbon- coated copper grids for 30 seconds. The grids were washed briefly with 0.1 and 0.01 M ammonium acetate buffer, pH 7.4, and stained with two 50 μL drops of freshly filtered 2% w/v uranyl acetate. The grids were allowed to dry overnight before viewing and the electron micrographs were recorded on FEI Eagle 4K bottom mount CCD camera. A minimum of three different preparations of each condition were examined by TEM, looking at least 5 different areas per grid.

### Mass spectrometry

#### Electrophoresis and In-Gel Protein Digestion

Samples from PTA-purified PrP^Sc^ and from SV and SE fractions showing HMW contaminating proteins >100 kDa were analysed by MS. 50 μL (of each sample was mixed with 16.6 μL of loading buffer (277.8 mM Tris-HCl, pH 6.8, 4.4% LDS, 44.4% (w/v) glycerol, 0.02% bromophenol blue, 10% (v/v) β-mercaptoethanol) and heated at 100°C for 10 minutes. 13 μL of each sample was resolved by SDS-PAGE (150 V for 60 minutes) using 10% sterile-filtered polyacrylamide gels, and subsequently fixed (50% (v/v) ethanol, 2% (w/v) phosphoric acid).

Gels were visualized with G-250 Coomassie Blue (SigmaAldrich) prior to whole-lane excision. Each lane was excised and cut into 15 equal bands, with each band further cut into 1 mm^3^ cubes. Gel fragments were destained (1:1 mixture of acetonitrile:100 mM ammonium bicarbonate), and subsequently reduced (10 mM DTT in 100 mM ammonium bicarbonate at 40°C for 1 hour) then alkylated (100 mM iodoacetamide in 100 mM ammonium bicarbonate at room temperature for 45 minutes) prior to in-gel trypsin digestion (30 μL of 6 ng/μL recombinant sequencing-grade trypsin (Promega) in 100 mM ammonium bicarbonate per cubed band) at 37°C overnight, as previously described [82].)The resulting tryptic peptides (per excised band) were subject to a three-step extraction (*i* – 1% (v/v) formic acid, 2% (v/v) acetonitrile in water; *ii* – 1% (v/v) formic acid, 50% (v/v) acetonitrile in water; *iii* – 1% (v/v) formic acid, 25% (v/v) water in acetonitrile), subsequently pooled, dried, and resuspended in 40μL of 1% (v/v) formic acid in 5% (v/v) acetonitrile.

#### Mass Spectrometry and Database Search Parameters

Individual fractions containing extracted peptides were resolved, ionized, and analyzed using nanoflow HPLC (Easy nLC-II, Thermo Scientific) coupled to a LTQ XL-Orbitrap hybrid mass spectrometer (Thermo Scientific). For nanoflow chromatography and electrospray ionization, a PicoFrit fused silica capillary column (ProteoPepII, C18; 100 μm inner diameter - 300Å, 5μm, New Objective) was utilized. Each peptide sample (5 μL) was subject to a 60 minute linear acetonitrile gradient (0% to 45% (v/v) acetonitrile in 0.2% (v/v) formic acid at 500 nL/minute).

Data-dependent acquisition mode was used to operate the LTQ XL-Orbitrap mass spectrometer, recording mass spectra using external mass calibration (full scan resolution of 60,000 and m/z range of 400–2000). The ten most intense multiply charged ions were sequentially fragmented with collision-induced dissociation, with fragment spectra recorded using the linear ion trap. Following two fragmentations, precursor ions selected for dissociation were subject to 60 second dynamic exclusion.

Raw data files corresponding to an entire gel lane were grouped and processed with Proteome Discoverer 1.4.1.14’s SEQUEST (Thermo Scientific) search algorithm using a reviewed, non-redundant Mus musculus complete proteome FASTA index (UniprotKB – retrieved October 2014). Search parameters were as follows: event detector mass precision – 2ppm; spectrum selector minimum and maximum precursor masses – 350 Da and 5000 Da, respectively, maximum collision energy – 1000; input data digestion enzyme – trypsin (full); maximum missed cleavage sites – 2; precursor mass tolerance – 10 ppm with 0.08 Da fragment mass tolerance; dynamic modifications to peptides – methionine oxidation (+15.995 Da), asparagine and glutamine deamidation (+0.984 Da); static modifications to peptides – cysteine carbamidomethylation (+57.021 Da). Relative extracted ion chromatograms for each identified protein were determined using the ‘Precursor Ion Area Detector’ node during data processing. Processed data was filtered using a minimum of 2 medium-confidence peptides per protein (FDR < 0.05), and exported to Microsoft Excel for analysis.

### Scrapie cell assay

All samples (whole dC2C12 lysates, PTA-purified PrP^Sc^ samples, and the fractions from SV) were methanol precipitated to remove detergents, iodixanol or other proteases or inhibitors (PMSF). Each fraction (200ul) was precipitated with 4 volumes of cold methanol for one hour and centrifuged at 16000xg for 30 minutes. The supernatant was carefully decanted off the pellet and the pellet air-dried. SSCA was performed as described previously [25], with some modifications. Briefly, L929 cells were exposed to the different PrP^Sc^ fractions resuspended in 100ul of culture medium; 30 µl was loaded per well, in triplicate, for 5 days in 96-well culture plates. The cells were passaged 3 times (1:4 and 1:7), with 20,000 cells collected at the third passage and loaded on to Multiscreen HTS IP 96-well, 0.45-μM filter plates (Millipore). The cells were PK digested (5 μg/ml) followed by denaturation using 3 M guanidine thiocyanate. The Elispot reaction was performed using a mouse anti-PrP antibody (SAF83, 1:1,000) and a goat anti-mouse alkaline phosphatase–conjugated secondary antibody (1:5,000). The plates were developed using BCIP/NBT and analyzed using an Autoimmun Diagnostika GmbH Elispot plate reader (ELR07).

### Immunohistochemistry for PrP and FN in paraffin tissues

Tissues collected from naturally scrapie-infected and control sheep were formalin-fixed and paraffin-embedded for IHC examination. Serial tissue sections (4 µm) assessed as previously described for PrP^Sc^ [10] and for FN detection. For PrP^Sc^ detection, briefly, sections were pre-treated with 98% formic acid, proteinase K (4 µg/ml; F. Hoffmann La Roche Ltd, 211 Switzerland) and by heating at 96°C for 20 min in EnVision^TM^ FLEX Target Retrieval solution. Then, tissue sections were incubated with a blocking reagent and immunolabeled using an automated immunostainer. Sections were incubated with an L42 primary antibody (0.046 µg/ml; R-Biopharm, Darmstadt, Germany) at room temperature for 30 minutes. EnVision was used as a visualization system and diaminobenzidine as the chromogen. For FN, sections were incubated with anti-FN pAb (Abcam, ab2413) using the protocol described above except only heating tissue sections in EnVision^TM^ FLEX Target Retrieval solution was performed. Finally, sections were counterstained with hematoxylin. Unless stated otherwise, all products were obtained from Dako (Denmark A/S, Denmark).

All tissues collected belong to a previous study Garza *et. al*., 2014 [10] approved by the Ethics Committee for Animal Experiments of the University of Zaragoza (Permit Number: PI02/08) and was carried out in strict accordance with the recommendations for the care and use of experimental animals and in agreement with the national legislation (R.D. 1201/2005).

## Acknowledgments

The authors gratefully acknowledge Pamela Banser for her help with the cell cultures.

## Supporting information

**S1 Fig. RML infected dC2C12 cell cultures at day 4 post inoculation observed by differential interference contrast (A, C) and phase contrast (B, D) microscopy.**

A fully differentiated C2C12 culture at day 4 post inoculation reveals the long myotubes in spiral patterns (A, B). Higher magnifications of multinucleated myotubes intermingled with mononucleated cells found at the upper layer of the culture (C, D). Scale bars 500µm (A, B), 100µm (C, D).

**S2 Fig. Histogram of myotube thickness in NBH-treated (white) and RML-infected (grey) dC2C12 culture.**

Thirty-eight myotubes from a minimum of 5 micrographs, at 40x magnification, were examined per culture and each value is the average of 10 measurements per myosegment; Chi squared equals 22.021 with 4 degrees of freedom. The two-tailed P value equals 0.0002; branched myotubes were not included in the analysis.

**S3 Fig. Immunoblot analysis of non-boiled and boiled (5, 15 and 30 min.) cell lysates after mild chemical cross-linking with 2% formaldehyde.**

A complex PrP-NCAM of 200 kDa is detected with both antibodies anti-PrP and anti-NCAM that disappear in the crosslink reversal time series.

**S4 Fig. Immunohistochemical detection of FN and PrP^Sc^ in heart and lymph node.**

No FN or PrP^Sc^ was detected in cardiac muscle (A,B). In lymph nodes (C,D), PrP^Sc^ was localized to the cortex of lymphoid follicles whereas FN was seen in the medulla associated with sinusoidal vessels. Sections A and C were incubated with anti-FN pAb (Abcam, ab2413) and sections B and D were incubated with L42 primary antibody (R-Biopharm, Darmstadt, Germany)

## Author Contributions

M Carmen Garza: **Conceptualization, Investigation, Visualization, Writing – Original Draft Preparation**

Sang Gyun Kang: **Conceptualization, Investigation, Visualization, Writing – Original Draft Preparation**

Leonardo Cortez: **Methodology, Supervision, Visualization, Writing – Original Draft Preparation**

Chiye Kim: **Methodology Investigation**

Eva Monleón: **Resources, Investigation**

Jacques van der Merwe: **Methodology**

David A. Kramer: **Resources, Investigation**

Richard Fahlman: **Resources, Investigation**

Valerie Sim: **Resources, Supervision, Review & Editing**

Judd Aiken: **Funding Acquisition, Supervision, Resources, Review and Editing**

Debbie McKenzie: **Funding Acquisition, Supervision, Resources, Review & Editing**

Holger Wille: **Funding Acquisition, Supervision, Review & Editing**

## Notes

### Competing Interest Statement

The authors have declared no competing interest.

